# Transposase-Assisted Tagmentation: An Economical and Scalable Strategy for Single-Worm Whole-Genome Sequencing

**DOI:** 10.1101/2023.12.21.572713

**Authors:** Zi Wang, Jinyi Ke, Zhengyang Guo, Yang Wang, Kexin Lei, Shimin Wang, Guanghan Chen, Zijie shen, Wei Li, Guangshuo Ou

## Abstract

AlphaMissense identifies 23 million human missense variants as likely pathogenic, but only 0.1% have been clinically classified. To experimentally validate these predictions, chemical mutagenesis presents a rapid, cost-effective method to produce billions of mutations in model organisms. However, the prohibitive costs and limitations in the throughput of whole-genome sequencing (WGS) technologies, crucial for variant identification, constrain its widespread application. Here, we introduce a Tn5 transposase-assisted tagmentation technique for conducting WGS in *C. elegans*, *E. coli*, *S. cervisiae*, and *C. reinhardtii*. This method, demands merely 20 minutes of hands-on time for a single worm or single-cell clones and incurs a cost below 10 US dollars. It effectively pinpoints causal mutations in mutants defective in cilia or neurotransmitter secretion and in mutants synthetically sterile with a variant analogous to the oncogenic BRAF(V600E) mutation. Integrated with chemical mutagenesis, our approach can generate and identify missense variants economically and efficiently, facilitating experimental investigations of missense variants in diverse species.

## Introduction

Utilizing Artificial Intelligence (AI), AlphaFold2 has formulated billions of protein structure models, elucidating protein structure-function relationships crucial to understanding organismal biology, while simultaneously advancing our knowledge of diseases and facilitating the development of novel therapeutics^1^. Recent advancements with AlphaMissense, a deep learning model built upon AlphaFold2, have categorized 89% of an estimated 71 million possible missense variants within the human proteome as either likely pathogenic or benign^2^. However, a mere 0.1% of such predictions have been corroborated by clinical data or functional studies^2^. Though CRISPR-Cas9-based genome editing strategies have been widely adopted to create genome-edited cell lines and animals to explore the implications of human missense mutations in model organisms^3–6^, the generation of single–amino acid substitutions across cell or animal models usually requires months of experimentation and involves costs upwards of several thousand US dollars, operating in a one-at-a-time manner. This approach contrasts starkly with the original aim of high-throughput loss-of-function or gain-of-function genetic screens that target the genome as a whole. Thus, there emerges a critical need to establish a rapid, economical, and scalable methodology to experimentally explore the physiological or pathological relevance of human missense variants.

Chemical mutagenesis, utilizing agents such as the alkylating compound Ethyl Methane Sulfonate (EMS) to alter DNA sequences and induce mutations, has been broadly employed across species in genetic and genomic research for decades^7–9^. For instance, Sydney Brenner pioneered the use of chemical mutagenesis in the model organism *Caenorhabditis elegans* (*C. elegans*) in the 1970s^10^. Adhering to the Brenner’s protocol, a single round of chemical mutagenesis typically yields approximately 91 missense variants among roughly 419 genomic alterations in an individual worm^11^. The simplicity of culturing nematodes facilitates the chemical mutagenesis of one million individual worms within just one week^9^, all at a nearly negligible cost. Since each mutagenized *C. elegans* produces about 100 progenies, chemical mutagenesis can thereby generate billions of missense variants both rapidly and economically^12^. The same holds true when chemical mutagenesis is applied to other model organisms, including bacteria^13,14^, yeast^15^, and algae^16^. Despite the potential and historical use of chemical mutagenesis, a major bottleneck impeding its widespread application is the financial burden associated with whole-genome sequencing (WGS) platforms, which are vital for identifying variants. A testament to this challenge is the Million Mutation Project (MMP), undertaken by the collective efforts of the community 12 years ago^11^. This project, which sequenced 2,007 chemically or UV mutagenized *C. elegans*, identified approximately 183,327 missense variants at a considerable expense of several million US dollars^11^. Despite its significance, the project was not continued, in part due to the then-prohibitive cost of WGS per strain, amounting to about 10,000 US dollars. Although the cost of WGS for a *C. elegans* strain has since diminished to several hundred US dollars, sequencing millions of strains remains economically unfeasible.

Beyond cost reduction, equally crucial for the application of chemical mutagenesis is minimizing the biological samples required to generate a WGS library. In a typical forward genetic screen of homozygous F2 *C. elegans*, only approximately 10% generate live F3 progenies with heritable phenotypes^9^. Even when exhibiting notable phenotypes, the remaining F2 mutant animals might, regrettably, be sterile or lethal that do not transmit to subsequent generations^9^. While theoretically plausible, maintaining heterozygotic mutants using a genetic balancer proves to be operationally tedious and is not universally applicable to every mutant. Thus, there is a significant demand to collect individual mutant worms to glean their genome information through WGS sequencing.

Bacterial transposase Tn5 is prevalently utilized in preparing various sequencing libraries due to its minimal sample input requirement and rapid processivity^17–20^. The Tn5 transposase dimer is distinctive for its unique tagmentation property: it can cleave double-stranded DNA (dsDNA)^17,18^ and concurrently ligate specific adaptors to the resultant DNA ends, a process subsequently followed by PCR amplification with sequencing adaptors^21,22^. This streamlined one-step tagmentation reaction has significantly simplified the experimental process, reducing both workflow duration and costs^21,22^. Tn5 tagmentation has been widely adopted for detecting chromatin accessibility and interactions^23–25^, as well as for enabling other types of sequencing experimentations^26^.

In this study, we present a Tn5 transposase-assisted tagmentation technique for conducting WGS of a single *C. elegans* specimen. This efficient protocol requires a mere 20 minutes of hands-on time and costs less than 10 US dollars. We show that the method is highly effective for pinpointing causal mutations in fertile mutants exhibiting defects in cilia or neurotransmitter secretion. Crucially, this method also facilitates the identification of mutations in mutants synthetically sterile with a variant analogous to the human BRAF(V600E) mutation. We demonstrate the applicability of this technique to one single clone of yeast or algae. This method holds great potential for widespread use in WGS of chemically mutagenized model organisms and possibly mammalian cell lines. When combined with chemical mutagenesis, our approach offers a cost-effective and efficient means of generating and identifying missense variants, thereby enhancing the experimental exploration of missense variants across a range of species.

## Results

### The single-worm WGS construction strategy

We introduce a rapid whole-genome sequencing (WGS) library construction method utilizing a single worm (Fig. 1A), comprising three primary components: *C. elegans* protein digestion, dsDNA tagmentation, and PCR amplification, culminating in an indexed library poised for sequencing. Initially, proteinase K is employed to digest a single worm. While an extended digestion duration (e.g., 15 hours) may facilitate comprehensive digestion of *C. elegans* proteins (Figure 1B-D, S1B), a concise two-hour digestion liberates sufficient DNA for subsequent experiments (Figure S1C-D). After the heat inactivation of proteinase K, the dsDNA undergoes tagmentation via the Tn5 transposase in the same tube, thereby appending partial sequencing adaptors to fragment ends. Thereafter, DNA polymerase amplifies the DNA fragments into a sequencing library. Amplified molecules were approximately 150 bp longer than the tagmentation products, corresponding to the additional length from the adaptors added during index primer amplification (Figure. S1A). This illustrates that Tn5 tagmentation of the *C. elegans* genomic DNA provides a feasible strategy for preparing a WGS library from an individual worm. The entire workflow requires just a single test tube and approximately 5 hours, with a hands-on time of under 30 minutes.

**Figure. 1.**
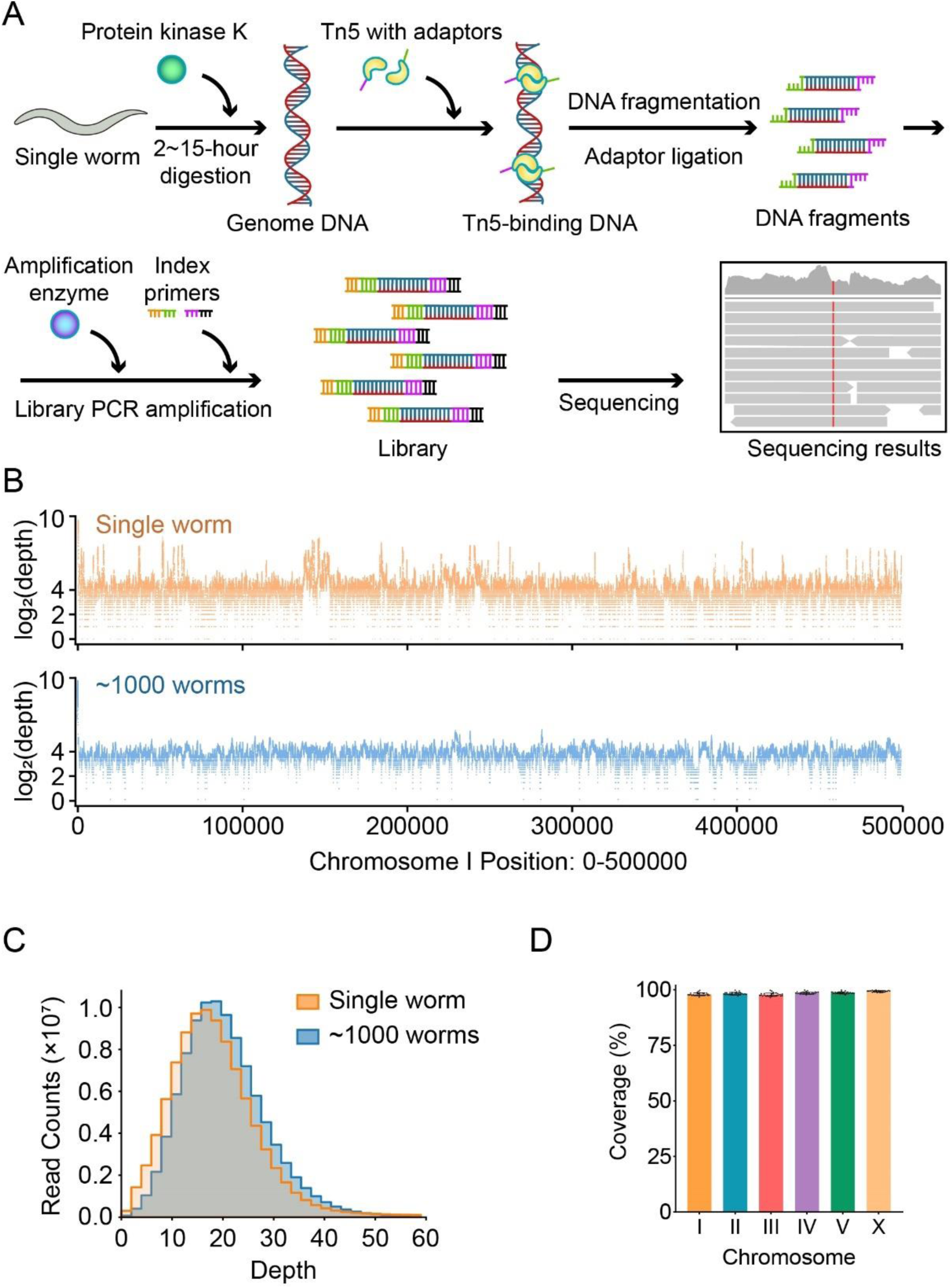
Whole-genome sequencing by direct tagmentation of a single worm. A. Scheme of single-worm WGS construction. B. The log_2_(depth) distribution along chromosome I position 0-500000 from a single worm or about 1000 worms. C. The read quantification of the depth from a single worm or about 1000 worms. D. Percentage of the coverage of each chromosome from a single worm with 15-hour PK digestion.

### The single-worm WGS results

In our analysis of the WGS data, we discovered that the sequencing results encompassed roughly 98% of the genome (Figure. 1D), with an average depth of 20× (Figure. 1B-C), a coverage that is comparable to existing WGS results obtained using approximately one thousand worms (Figure S2B for comparison). Extending our initial success to an additional 29 individual worms, we achieved highly reproducible WGS results (Figure. S1B). Furthermore, when we plotted the WGS coverage data for all 29 worms on each chromosome, we observed consistently high, unbiased coverage, further attesting to the robustness of the method (Figure. 1D).

### The single-worm WGS identified mutations responsible for mutant phenotypes

We explored whether the WGS results obtained from a single worm would enable the identification of genetic mutations induced by EMS mutagenesis. *C. elegans* utilize their sensory cilia to engage with environmental stimuli^27,28^. In wild-type worms, which develop normal cilia, a carbocyanine dye, DiI, can be absorbed through sensory cilia^28^. Conversely, cilia mutants, which fail to fill with DiI, exhibit a dye-filling defect (Dyf) phenotype, making the dye filling assay a powerful tool for isolating ciliary mutants^28^. We found that the *cas2885* strain did not uptake DiI, and our WGS results from a single mutant animal revealed that *cas2885* harbors a newly acquired stop codon in *dyf-5* (Figure 2A), which encodes a ciliary kinase essential for ciliary length regulation^29,30^. To validate that this mutation is responsible for the Dyf phenotype, we injected the WT *dyf-5* cDNA under the control of the ciliated neuron-specific promoter, P*dyf-1*, into the *cas2885* strain. We demonstrated successful restoration of the Dyf defects in two independent transgenic lines (Figure 2B).

**Figure. 2.**
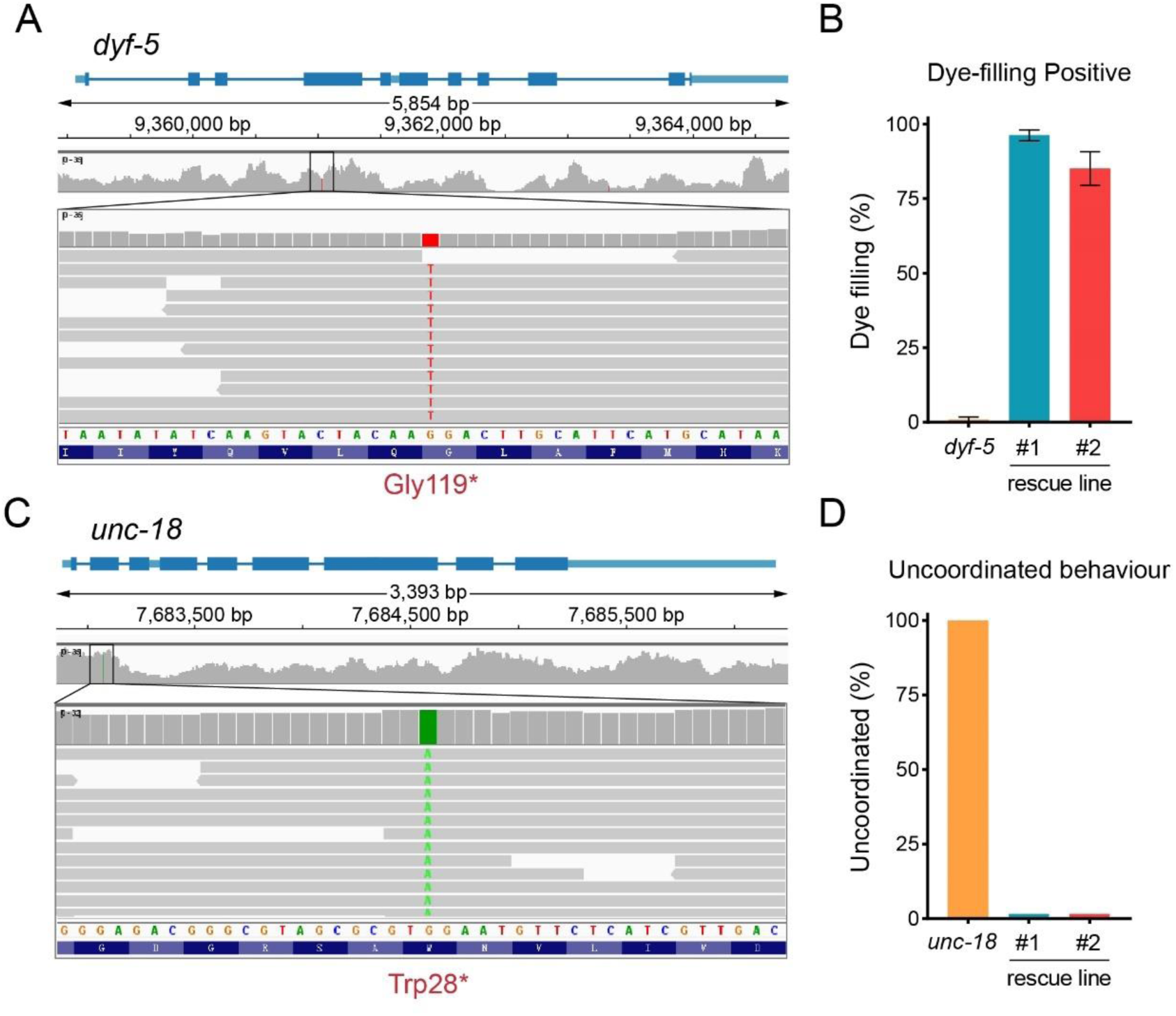
The single-worm WGS identified mutations responsible for mutant phenotypes. A. An IGV window shows the mutation site of the *dyf-5* mutant. B. Percentages of Dye-filling positive *dyf-5* mutant and the *dyf-5* rescue worms. n > 50. C. An IGV window shows the mutation site of the *unc-18* mutant. D. Percentages of uncoordinated behavior *unc-18* mutant and the *unc-18* rescue worms. n > 50.

Similarly, we employed EMS mutagenesis to generate a cohort of animals exhibiting uncoordinated (Unc) movement. Among them, our single-worm WGS identified a point mutation leading to an ectopic stop codon in the *unc-18* gene (Figure 2C), which regulates neurotransmitter secretion^31^. By introducing the wild-type *unc-18* gene into the *cas4401* strain carrying this mutation, we observed a complete rescue of its Unc phenotype in two independent transgenic lines (Figure 2D). Thus, the examples of *dyf-5* and *unc-18* validate that the single-worm WGS results enable us to identify genetic variants responsible for animal phenotypes.

### The single-worm WGS identified mutations from a sterile mutant

We investigated the potential of our method in single worms that are progeny-deficient. Specifically, we performed a genetic enhancer screen on a strain with a LIN-45 missense mutation V627E, analogous to the pathogenic BRAF(V600E) mutation in humans (Figure 3A). The *BRAF* gene, a critical component of the RAS/RAF/MEK/ERK signaling pathway, is pivotal in regulating cell division and differentiation^32–34^. LIN-45 serves as the *C. elegans* ortholog of BRAF^35^. The V600E mutation in BRAF leads to excessive activation of the kinase, triggering uncontrolled cell division and contributing to cancer development^32,36–39^. Employing genome editing, we created a *C. elegans* strain with the corresponding *lin-45(V627E)* mutation. This mutation manifested as a protruding vulva phenotype with ectopic cells due to overproliferation.

**Figure. 3.**
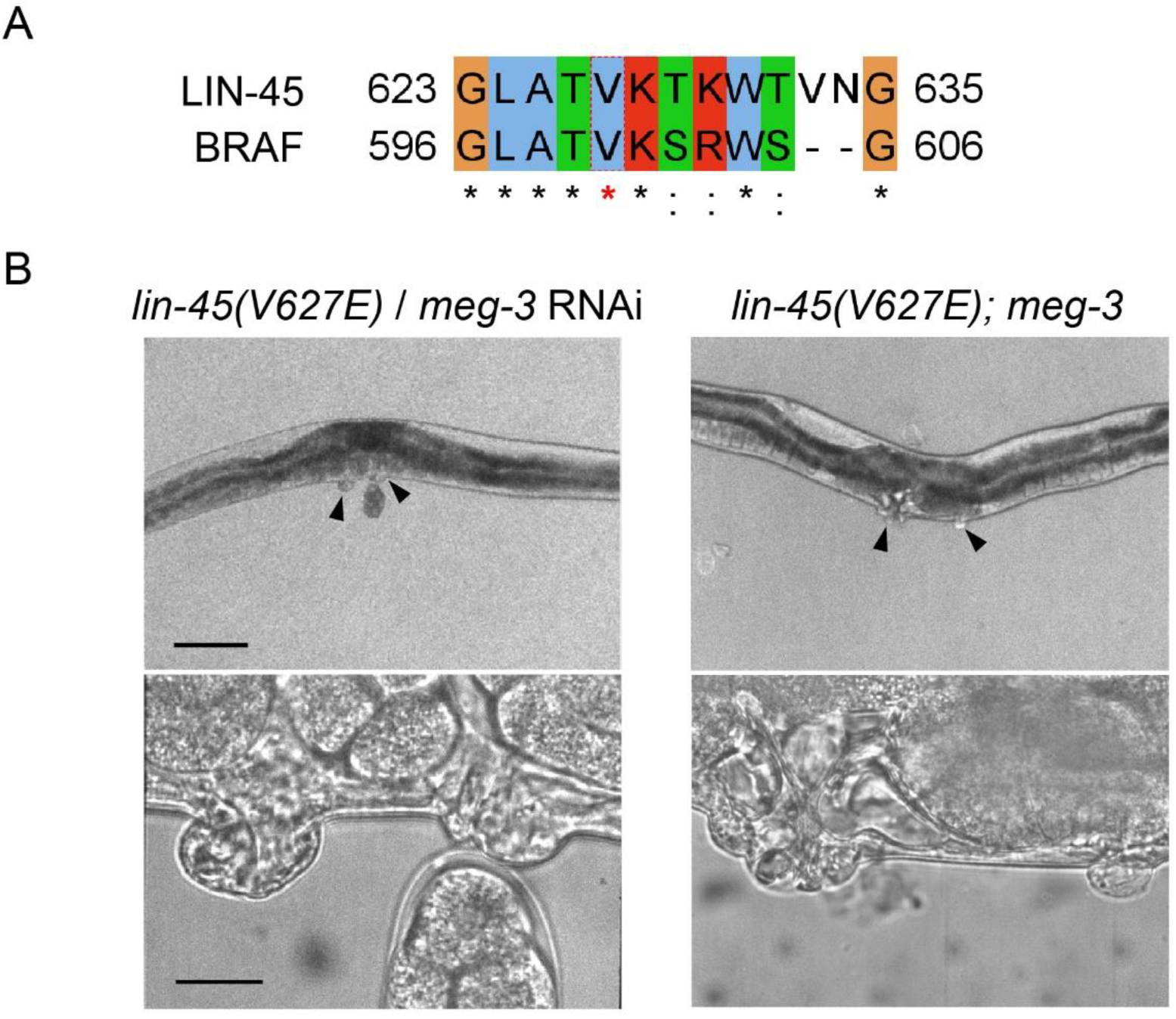
The single-worm WGS identified mutations from a sterile mutant. A. An Jalview window shows a part of the alignment between the human BRAF and *C. elegans* LIN-45. B. Representative images of the vulva of *lin-45(V627E)* treated with *meg-3* RNAi and *lin-45(V627E); meg-3*. For 10× magnification images, scale bar is 100 μm. For 100× magnification images, scale bar is 10 μm.

During our enhancer screen across ∼20,000 haploid genomes, we isolated 30 suppressors from the F2 generation displaying a multi-vulva phenotype, likely due to additional unregulated cell divisions. Remarkably, only nine of these suppressors produced viable progeny exhibiting an intensified vulva phenotype; the rest were sterile. We conducted single-worm Whole Genome Sequencing (WGS) on these sterile lines and discovered a stop-gain mutation in the *meg-3* gene, a known regulator of cell fate (REF)^40^. When *meg-3* was knocked down via RNAi in the LIN-45 V627E strain, about 4.8% of the 96 progenies showed the multivulva phenotype (Figure 3B). While the penetrance was low, this phenotype was absent in *meg-3* RNAi-treated wild-type animals and in the negative control RNAi *lin-45(V627E)* strain. To further examine this, we used CRISPR-Cas9 to generate *meg-3* knockout strains in the *lin-45(V627E)* background. We found that *meg-3* null mutants with the *lin-45(V627E)* strain developed an enhanced vulva phenotype (Figure 3B) but were unable to produce offspring. These results demonstrate how single-worm WGS can effectively pinpoint causative mutations in sterile strains.

### Effective single-clone WGS in bacteria, yeast and algae

To evaluate the applicability of our method in other species, we wondered its efficacy in performing WGS on single clones of *E. coli*, *S. cervisiae*, and *C. reinhardtii*, each comprising approximately 1000 cells—comparable to the 959 somatic cells of an individual *C. elegans*. The identical protocol adeptly generated the WGS library using the cell number equivalent to ones from a single clone, the sequencing of which yielded a coverage rate and depth analogous to that of a single *C. elegans* (Figure 4A-C, S3, S4).

**Figure. 4.**
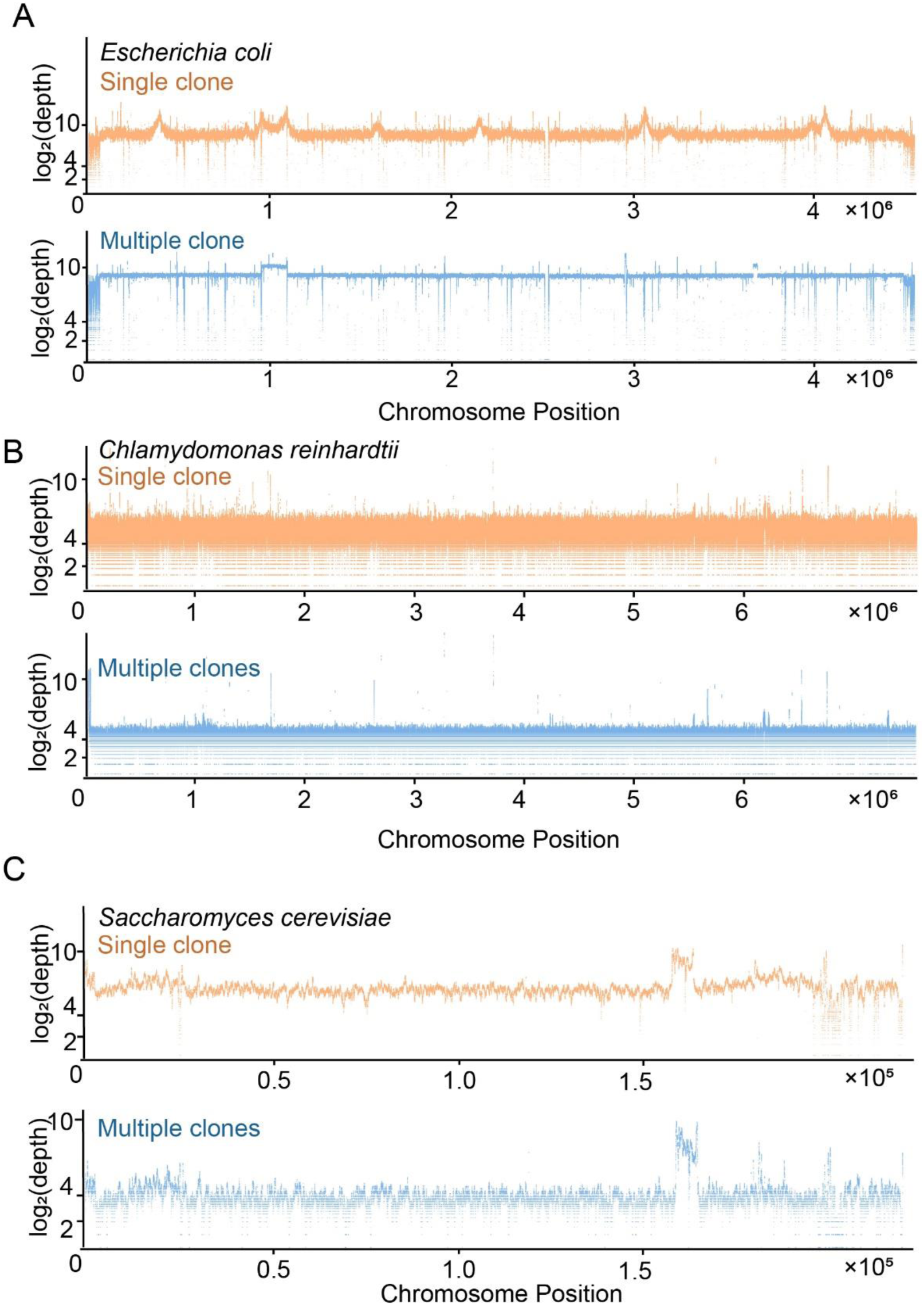
Effective single-clone WGS in bacteria, yeast and algae. A. The log_2_(depth) distribution along the chromosome from a single clone or multiple clones of *Escherichia coli*. B. The log_2_(depth) distribution along chromosome I from a single clone or multiple clones of *Chlamydomonas reinhardtii*. C. The log_2_(depth) distribution along chromosome I from a single clone or multiple clones of *Saccharomyces cerevisiae*.

## Discussion

In summary, our findings demonstrate that Tn5 transposase-assisted tagmentation can facilitate the development of a methodology capable of generating DNA libraries for whole-genome sequencing from a single worm. The same method is applicable across other species, such as a single clone of yeast or algae. The implementation of the protocol is notably straightforward. The single-worm PCR protocol, frequently utilized for *C. elegans* genotyping, is often considered an entry-level experiment for novice researchers in *C. elegans* research groups or even in educational laboratories^41,42^. Our single-worm WGS protocol merely introduces an additional Tn5 reaction step in the same tube, underscoring its operational simplicity. As outlined in the Materials and Methods section, the single-worm WGS library construction is accomplished within a 15-microliter reaction volume, with reagent costs amounting to only $3. Given that the *C. elegans* genome is approximately 100 Mb in size, generating 2 Gb of data for a 20× coverage rate comes at a minimal cost of $7. Notably, many labs opt to sequence the genome at a 5× coverage rate, effectively halving the sequencing cost with 1 Gb of data, reducing the total to $10 or even less. Considering that all reactions can be conducted in a tube within a 96-well plate, after a single worm or clone is placed into the tube, all subsequent steps can potentially be automated. This not only minimizes experimental errors but also enhances scalability. Given that Tn5 transposases also act on DNA/RNA hybrids, protocols for constructing an RNA-seq library have been established in previous studies^19,20^. Consequently, we posit that generating both WGS and RNA-seq libraries from a single worm is plausible, thereby potentially furnishing both genomic and transcriptomic information from an individual *C. elegans*.

While the per-missense variant generation in *C. elegans* is estimated to be mere cents, it is imperative to note that this production strategy leans on random mutagenesis, not on a precision-targeted knock-in substitution. The probability of obtaining a desired missense variant is tethered to the size of the worm library fortified with WGS information. On average, *C. elegans* proteins encompass about 470 amino acids, and with its genome harbors approximately 20,000 protein-encoding genes^43^, this culminates in around 9.4 million residues within the *C. elegans* proteome. Given that each chemically mutagenized worm typically carries around 91 missense mutations^11^, conducting WGS on 0.1 million *C. elegans* mutant strains could, in probability, mutate each residue once. This results in a single-fold mutation coverage at an estimated expenditure of one million US dollars. Although EMS mutagenesis typically favors GC to AT conversions^44^, the incorporation of an additional mutagen, ENU^45^, is frequently employed to formulate “cocktail” mutagens^11^, thereby augmenting the diversity of mutation forms. Hence, with the allocation of additional resources, achieving a comprehensive coverage of various types of missense variants becomes plausible. While this protocol is established in simpler model organisms, we anticipate its broad application across numerous human cell lines, including those with a haploid genome that are primed for chemical mutagenesis-based forward genetic screens.

The single-worm/clone WGS methodology stands poised to expedite the functional study of missense variants identified within the human proteome, thereby harboring the potential to advance precision genomic medicine to a nucleotide-resolution tier. Alterations to individual amino acid residues within a protein may lead to distinct dysfunctions at biochemical, cellular, and organismal levels, each likely demanding unique interventional approach. This predicament underscores an emerging field, termed ’functional residuomics’, which endeavors to provide residue-resolution functional insights into the proteomic landscape.

The generation of missense variants in model organisms constitutes a foundational step in functional residuomic studies. Organisms harboring missense variants swiftly provide empirical evidence, crucial for differentiating benign from pathogenic variants. Characterizing the cellular and organismal impacts of pathogenic variants enables the acquisition of invaluable mechanistic insights into the interplay between genetic anomalies and symptomatic manifestations. Crucially, organisms that carry pathogenic variants present a starting point for executing genetic suppressor screens. This can illuminate strategies for phenotype rescue, potentially paving the way for effective clinical interventions and informing drug discovery endeavors.

If millions of strains are sequenced via WGS, a paramount challenge emerges in the storage and distribution of the sequenced strains. Presently, such reagents are deposited into genetic centers, like the *C. elegans* Genetic Center, which has been distributing strains for over four decades. Nevertheless, non-profit resource centers, bounded by limited government support, cannot expand indefinitely, a constraint equally applicable to commercial services like Addgene that distributes published plasmids. No institute akin to these centers possesses the capacity to store and distribute millions of strains with distinct genetic backgrounds, especially as these numbers perpetually augment. A de-centralization mechanism, reminiscent of eBay, may be a forward-thinking solution to navigate this predicament: each laboratory conducts their WGS and data analysis, depositing the sequencing information into a public platform or database. While a lab might concentrate on a missense mutation of interest to their work, others might scour the database for additional valuable variants. This platform not only supports the exchange of WGS information but also fosters the trade of strains, allowing laboratories to negotiate expenses, thereby encouraging a collaborative and mutually beneficial scientific environment.

## Materials and Methods

### Strains and Genetics

*C. elegans* strains were maintained as described previously^10^, on nematode growth medium (NGM) plates (3 g/l NaCl, 17 g/l agar, 2.5 g/l peptone, 1 mM CaCl_2_, 1 mM MgSO_4_, 25 mM KPO_4_ / pH 6) with OP50 feeder bacteria at 20°C. All the engineered *C. elegans* strains were genetic derivatives of the strain Bristol N2. Strains used in this study is summarized in ***Table S1***. Transformation of *C. elegans* to introduce the P*dyf-1::dyf-5* and P*unc-18::unc-18* rescue strains were performed by DNA injection as described^46^, and the information of plasmids and primers are described in ***Table S2.*** All animal experiments were performed following governmental and institutional guidelines. For OP50 culturation, one clone of OP50 was streaked onto LB Agar plate (15 g/l agar, 10 g/l tryptone, 5 g/l yeast extract, 10 g/l NaCl) and incubated overnight at 37°C.

Motile *Chlamydomonas reinhardtii* wild-type strain 21gr were grown in standard Tris-acetate phosphate medium (2.42g Tris, 1×TAP salt, 0.114g K_2_HPO_4_, 0.054 g KH_2_PO_4_, 0.1% Hutner’s trace metals, 0.1% glacial acetic acid, 0.4 g NH_4_Cl, 0.05 g CaCl_2_·2H_2_O, 0.1 g MgSO_4_·7H_2_O) in cycles of 12 h in fluorescent white light and 12 in darkness at 200 rpm at 20°C.

### Transposase-Assisted Tagmentation and Library Construction

For single-worm WGS, a solitary *Caenorhabditis elegans* specimen was employed as the experimental unit. In the case of single clones of *Escherichia coli*, *Saccharomyces cerevisiae*, or *Chlamydomonas reinhardtii*, each microorganism was initially isolated and suspended in PBS buffer (Sigma, 806544). Subsequently, the isolated clones were enumerated using a cell counting board under a dissecting microscope, with approximately 1000 cells selected as representative samples within each experimental group. Samples were placed into separate PCR tubes containing 3 μl of lysis buffer with 0.3 mg/ml protein kinase K (Thermofisher, 26160). Subsequently, freezing in liquid nitrogen for 1 minute, followed by thawing in a 37°C water bath for 2 minutes, was performed for 3 times to lyse the cell protein. Afterward, the samples were subjected to a specific program in a PCR instrument, involving incubation at 65°C for 2 or 15 hours, 95°C for 15 minutes, and a 4°C hold. The resulting DNA samples underwent fragmentation by mixing with thawed reagents, including 1 µl 5×TTBL and 1.25 µl TTE Mix (Vazyme, TD502/TD503), and were then treated at 55°C for 10 minutes in a PCR instrument. Following this, 1.25 µl of 5×TS (Vazyme, TD502/TD503) was added, and the mixture was incubated at room temperature for 5 minutes. Subsequently, 2.5 µl of adaptors mix (Vazyme, TD202), 0.25 µl of amplification enzyme (Vazyme, TD502/TD503) and 1 µl of double distilled water (ddH_2_O) were added to the PCR tube on ice, followed by amplification through a PCR program with defined temperature cycles (Vazyme, TD502/TD503). Finally, amplified libraries each with individual adaptor pairs, were purified using the PureLink Quick PCR Purification Kit (Invitrogen, #00995126).

### Forward genetic screens

We used forward genetic screens to isolate dye-filling defective animals, uncoordinated animals and multivulva animals. Adult animals were bleached by bleach buffer (1.26% NaHClO, 0.25 M NaOH) for 1.5 minutes to lyze worms and get eggs, which were then washed by M9 buffer (5.8 g/l Na_2_HPO4, 3.0 g/l KH_2_PO4, 0.5 g/l NaCl, 1.0 g/l NH_4_Cl) for 3 times and hatched on NGM plates with fresh OP50. Worms were used as P0 and collected at the late L4 larval stage in 4 mL M9 buffer, and incubated with 180 mM ethyl methane sulfonate (EMS) for 4 hours at room temperature with constant rotation. Animals were then washed with M9 three times and cultured under standard conditions. After 20 hours, adult animals were bleached. Eggs (F1) were distributed and raised on ∼100 9-cm NGM plates, each containing 50 to 100 eggs. For the dye-filling defective animals, *osm-3::gfp(syd0199)* was used as P0. Adult F2 animals on each plate were collected and examined via the dye-filling assay (see Dye-filling assay below). For the uncoordinated animals, *osm-3::gfp(syd0199)* was used as P0. F2 animals with uncoordinated phenotype were collected. For multivulva animals, the *lin-45 (syb4962)* animals were used as P0. F2 animals with multivulva phenotype were collected. We identified mutations using whole-genome sequencing. We confirmed gene cloning using rescue experiments.

### Microinjection and transgenic strains

Tran*sgenic or knockout C. elegans lines were generated by injecting the DNA constructs* (Table S3) into the gonads of the indicated worm strains. The co-injected selection marker was *pRF4(rol-6)*. At least two independent transgenic lines with a constant transmission rate (>50%) were propagated. Concentrations of DNA constructs used for generating rescue strains was 20 ng/μl.

### Dye-filling assay

The fluorescence dye DiI filling assay was widely used to assess the ciliary function and integrity. Animals that are dye-filling defective develop abnormal ciliary structures and are defective in animal behavioral assays, such as the osmotic avoidance assay and chemotaxis assay. Young adult worms were harvested into 100∼200 μL M9 buffer and mixed with equal volume dyes (DiI, 1,1’-dioctadecyl-3,3,3’,3’,-tetramethylindo-carbocyanine perchlorate, Sigma, 468495) at the working concentration (20 μg/ml), followed by incubation at room temperature in the dark for 30 min. Worms were then transferred to seeded NGM plates and examined for dye uptake two hours later using a fluorescence stereoscope or fluorescence compound microscope. We observed at least 50 animals of each strain from three independent assays.

### RNAi by feeding

Young adult C. elegans hermaphrodites were anesthetized with 0.1 mM/L levamisole (Sigma, Y0000047) in M9 buffer, mounted on 3% agarose pads, and maintained at room temperature. Imaging was performed using a Zeiss Axio Observer Z1 microscope equipped

### Imaging

Young adult C. elegans hermaphrodites were anesthetized with 0.1 mM/L levamisole (Sigma, Y0000047) in M9 buffer, mounted on 3% agarose pads, and maintained at room temperature. Imaging was performed using a Zeiss Axio Observer Z1 microscope equipped with an Andor iXon+ EM-CCD camera, a Zeiss 10×/0.45 objective, and a Zeiss 100×/1.46 objective. Images were acquired by µManager (https://www.micro-manager.org). All the images were taken using identical settings. Image analysis and measurement were performed with ImageJ software (http://rsbweb.nih.gov/ij/).

### Next generation sequencing (NGS) and data analysis

After library preparation, the DNA libraries were subjected to an Illumina Nova seq for Paired end 150bp whole genome sequencing (WGS) (Table S4 for detailed strain and raw base count). Raw reads were assessed for quality with FastQC (version 0.11.9) and were trimmed using Trim_galore (version 0.4.4) to remove the adaptor sequence and low-quality reads. After that, clean reads were aligned to the reference genomes using BWA-MEM2 (version 2.2) with default parameters. PCR duplications were marked and removed with Picard (2.27.5 and OpenJDK 20.0.2) MarkDuplicates tool. Information about the depth and coverage of sequencing were generated by samtools and further analyzed by a python script to generate the whole-genome scale coverage and depth plots. Reference genome of every organism sequenced were listed in ***Table S5***, except E. coli, the sequence result of all the other organisms were compared to their up-to-date standard reference genomes. all the shell and python scripts used in the paper are available in the github repository: young55775/single-worm-sequencing (github.com)

## Acknowledgments

The inception of the tagmentation-based methodology for single-worm whole-genome sequencing germinated from deliberations with Professors Yanyi Huang, Jianbin Wang, and Di Chen during the 2023 Artificial Evolution Summer School, which was generously supported by Tsinghua University and Qinshan Lake Science and Technology City, located in Hangzhou, China. Motile *Chlamydomonas reinhardtii* wild-type strain 21gr was kindly provided by Professor Junmin Pan. NYM5 was a gift from Professor Shanjin Huang. This work was supported by the following funding programs: the National Natural Science Foundation of China (grants 32270721, 31991190, 32070706, 32021002, 31970180, 31900535, 32071191 and 31971160), the National Key R & D Program of China (2017YFA0503501, 2019YFA0508401, and 2017YFA0102900).

---

**Figure. S1.**
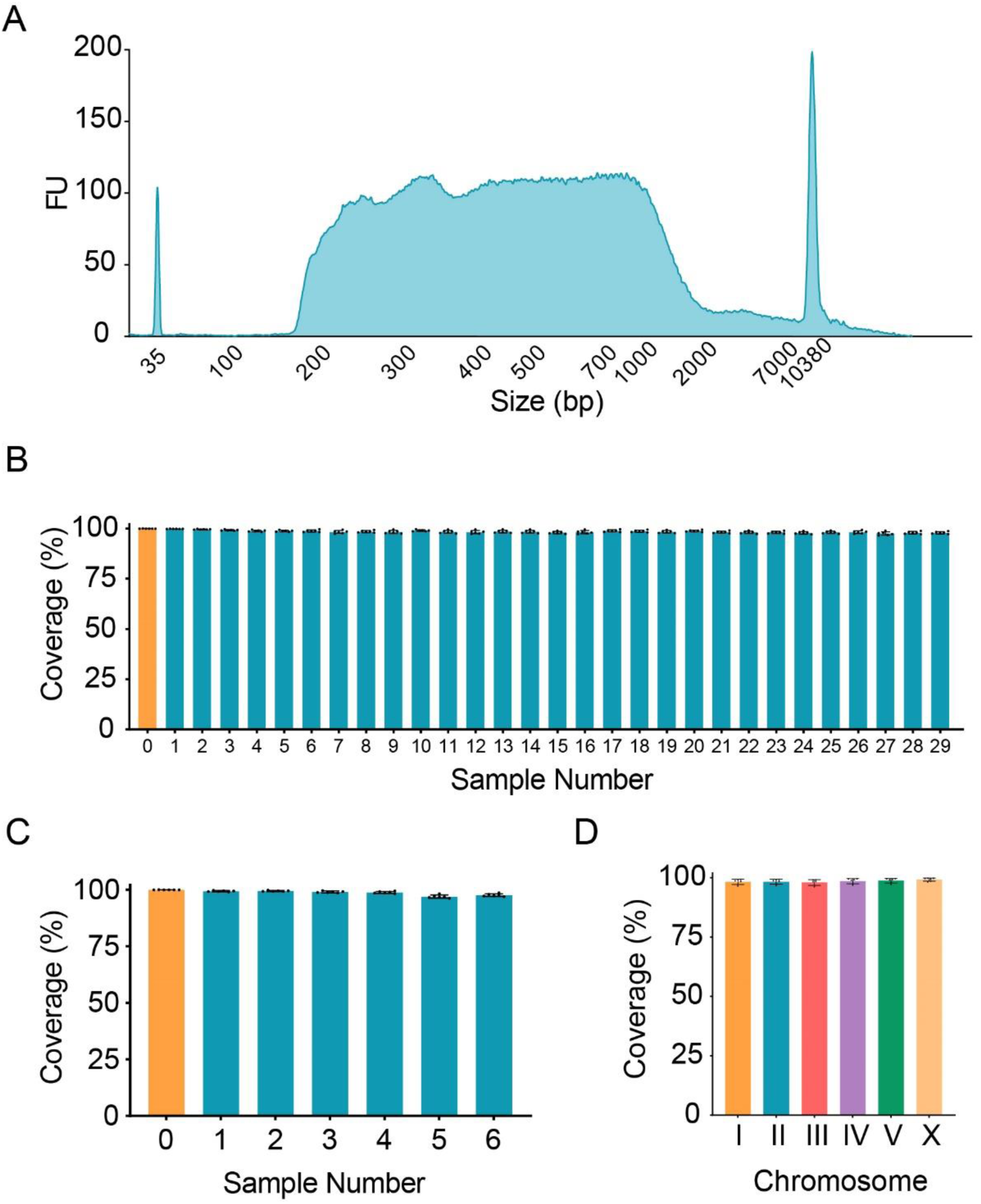
Quality of single-worm WGS. A. Representative Bioanalyzer electropherogram of a single-worm whole-genome library. B-C. Percentage of the coverage of each chromosome from individual worm sequencing data. B. The orange bar represents one traditional WGS data, blue bars represent 29 single-worm WGS data from worms with 15-hour PK digestion. C. The orange bar represents one traditional WGS data, blue bars represent 29 single-worm WGS data from worms with 2-hour PK digestion. D. Percentage of the coverage of each chromosome from a single worm with 2-hour PK digestion.

**Figure. S2.**
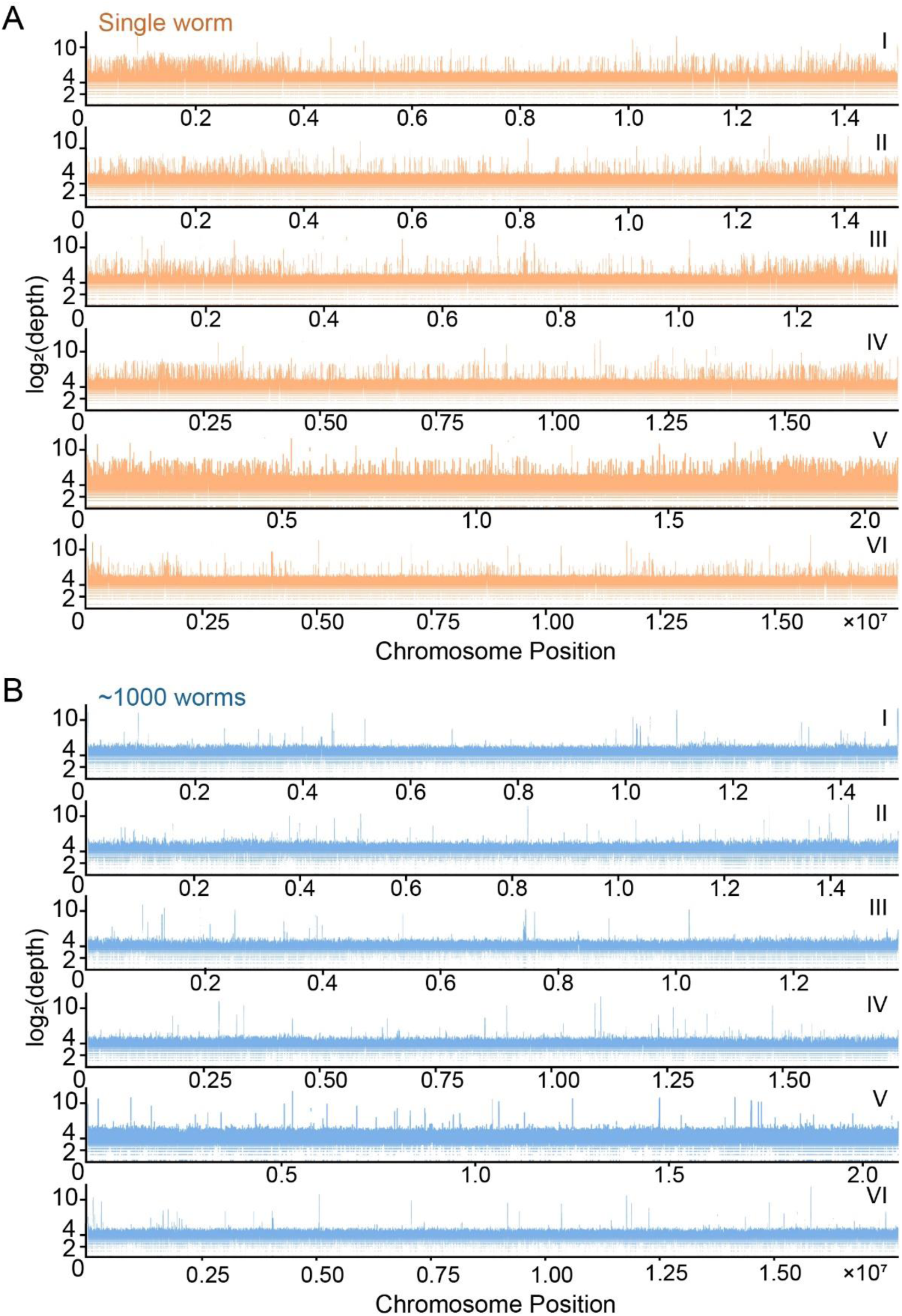
Quality comparison between single-worm WGS and traditional WGS. A. The log_2_(depth) distribution along individual chromosomes from a single *Caenorhabditis elegans*. B. The log_2_(depth) distribution along individual chromosomes from about 1000 *Caenorhabditis elegans*.

**Figure. S3.**
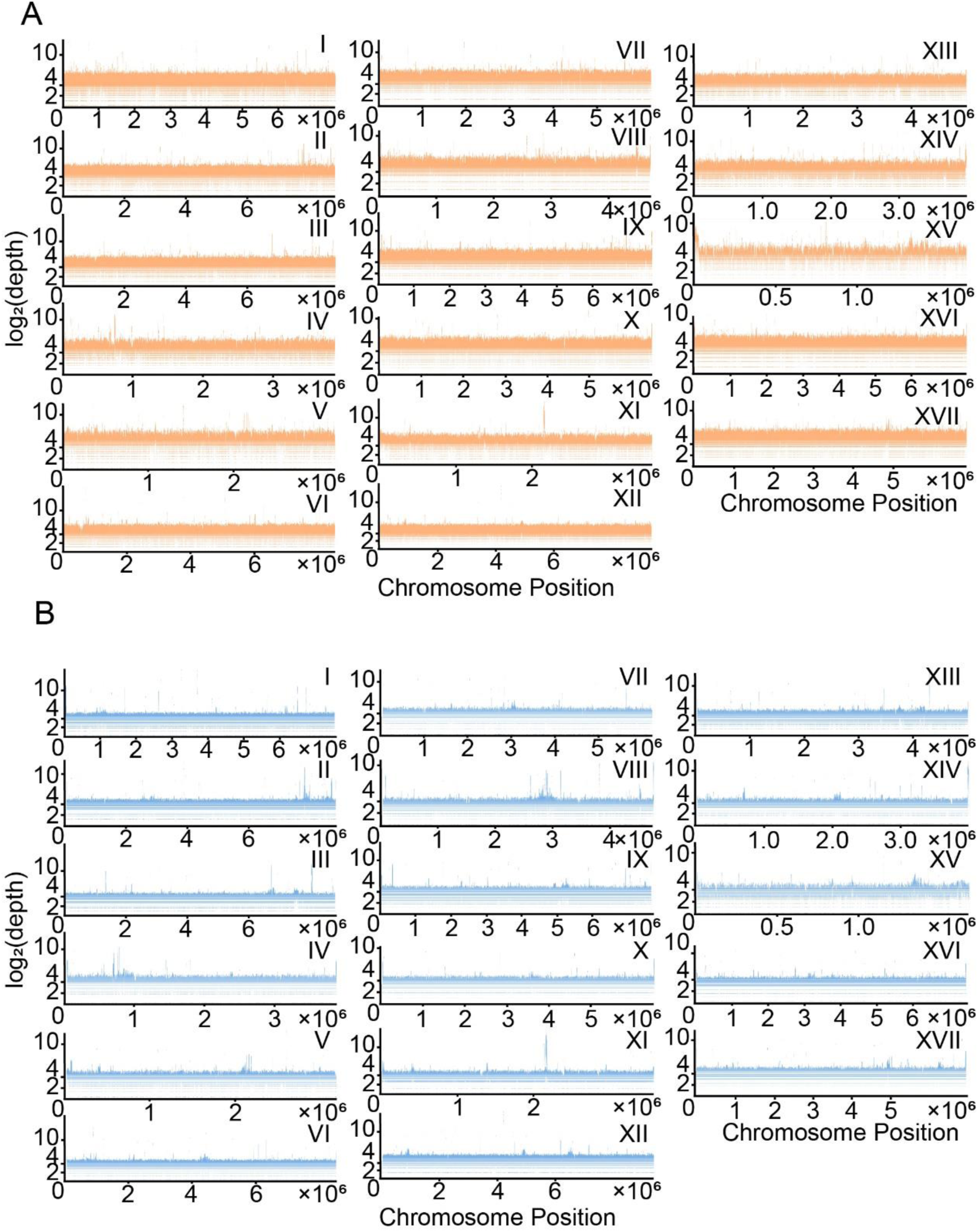
Quality comparison between single-clone WGS and traditional WGS in *Chlamydomonas reinhardtii*. A. The log_2_(depth) distribution along individual chromosomes from a single clone of *Chlamydomonas reinhardtii*. B. The log_2_(depth) distribution along individual chromosomes from multiple *Chlamydomonas reinhardtii*.

**Figure. S4.**
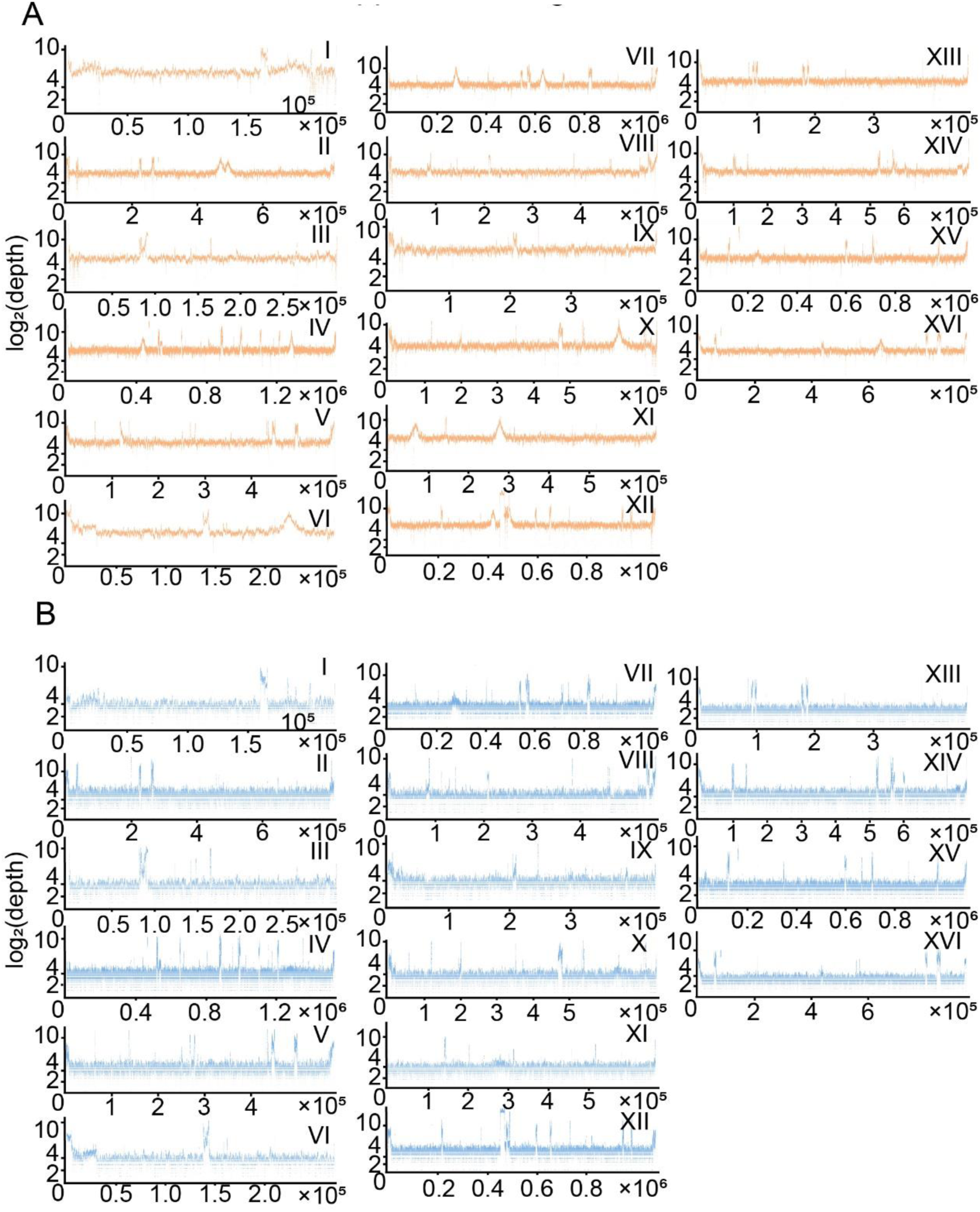
Quality comparison between single-clone WGS and traditional WGS in *Saccharomyces cerevisiae*. A. The log_2_(depth) distribution along individual chromosomes from a single clone of *Saccharomyces cerevisiae*. B. The log_2_(depth) distribution along individual chromosomes from multiple *Saccharomyces cerevisiae*.

**Table S1.** *C. elegans* Strains in this study.

| Strain Name | Genotype | Method |
| --- | --- | --- |
| SYD0199 | *osm-3(syd0199[osm-3::gfp KI])* |  |
| *cas2885* | *dyf-5(G119\*); osm-3(syd0199[osm-3::gfp KI])* | EMS screen |
| *Cas4401* | *unc-18(W28\*); osm-3(syd0199[osm-3::gfp KI])* | EMS screen |
| *cas2940* | *dyf-5(G119\*); osm-3(syd0199); Ex[dyf-1p::dyf-5 + rol-6(su1006)]* line 1 | Microinjection |
| *cas2941* | *dyf-5(G119\*); osm-3(syd0199); Ex[dyf-1p::dyf-5 + rol-6(su1006)]* line 2 | Microinjection |
| *cas2942* | *unc-18(W28\*); osm-3(syd0199); Ex[unc-18p::unc-18 + rol-6(su1006)]* line 1 | Microinjection |
| *cas2943* | *unc-18(W28\*); osm-3(syd0199); Ex[unc-18p::unc-18 + rol-6(su1006)]* line 2 | Microinjection |

**Table S2.**
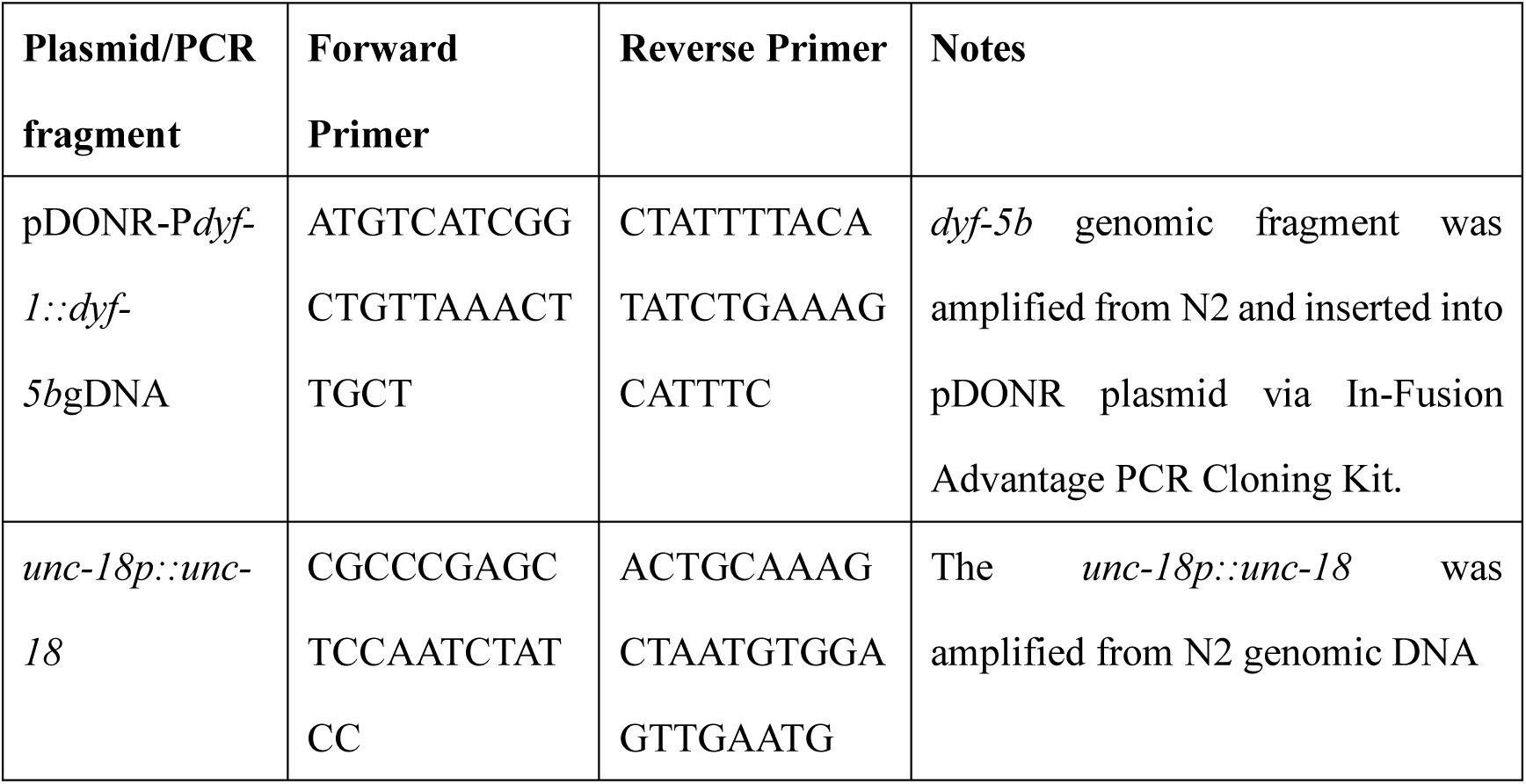
Plasmids, PCR fragments, and Primers in this study.

**Table S3.**
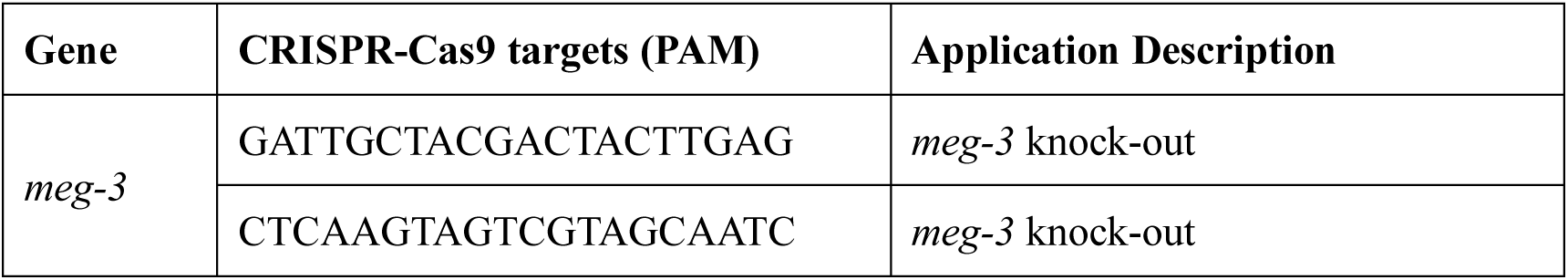
CRISPR-Cas9 Targets in this study.

**Table S4.** Species for sequencing.

| **Taxonomy** | **strain** | **Reference genome** |
| --- | --- | --- |
| *Caenorhabditis elegans* | N2 | Caenorhabditis_elegans – Ensembl genome browser 110 |
| *Saccharomyces cerevisiae* | NYM51 | Saccharomyces_cerevisiae - Ensembl genome browser 110 |
| *Escherichia coli* | OP50 | https://www.ncbi.nlm.nih.gov/datasets/genome/GCF_013456775.1/ |
| *Chlamydomonas reinhardtii* | 21gr | Chlamydomonas_reinhardtii - Ensembl Genomes 57 |

**Table S5.** Softwares for sequencing analysis.

| **software** | **version** | **link** |
| --- | --- | --- |
| FastQC | 0.11.9 | https://github.com/s-andrews/FastQC |
| Trim_galore | 0.4.4 | github.com |
| BWA-MEM2 | 2.2 | github.com |
| Picard | 2.27.5 | broadinstitute/picard: A set of command line tools (in Java) for manipulating high-throughput sequencing (HTS) data and formats such as SAM/BAM/CRAM and VCF. (github.com) |
| samtools | 1.18 | samtools/samtools: Tools (written in C using htslib) for manipulating next-generation sequencing data (github.com) |

